# BC-store: a program for mgiseq barcode sets analysis

**DOI:** 10.1101/2020.10.28.355156

**Authors:** Irina Bulusheva, Vera Belova, Boris Nikashin, Dmitriy Korostin

## Abstract

Here we present the devised BC-store – a program for analyzing and selecting sets of barcodes for sequencing on platforms manufactured by MGI Tech (China). The app is available as an open source in Python3 and as a desktop version. The application allows analyzing the compatibility of barcodes on a single lane of a flow cell in a set in the case of equal and arbitrary fractions. In addition, with the help of this tool barcodes can be added to an existing set with custom share options. In this paper we describe how BC-store works for different tasks and consider the effectiveness of using BC-store in sequence lab routine tasks.

**Author summary:** The performance of modern NGS machines allows considerable amount of data to be obtained which exceed the data required for one specific sample. To pool multiple samples on a single lane of the flow cell, barcoding is used – adapters carrying a unique nucleotide sequence are introduced by ligation [1]. Adapters are sequenced from their specific primers. Their sequences are used for demultiplexing the sequencing results for individual fastq files by programs like zebracall [2] or bcl2fastq [3]. The task of selecting the adapters for a set is similar to sequencing low diversity libraries [4]: if all adapters on the lane have the same nucleotide during this sequencing cycle, the quality of its reading drops dramatically. Therefore, manufacturers recommend grouping the adapters by sets. However, the sets offered by MGI Tech [5] are far from routine practice, as they do not allow for non-equimolar sample pooling by default, and they have other disadvantages (more on them later in the text). To overcome these problems, we created the BC-store program, which allows analyzing the sets of MGI Tech barcodes entered by the user to be further used in sequencing. This tool also provides the opportunity to vary the number of simultaneously sequenced samples, to use reagents more efficiently, and, as a result, to correctly distribute reads across samples, which helps to increase both the quality of sequencing and the level of data interpretation.

## Introduction

The barcode for MGISEQ-2000/DNBseq-G400 is a sequence of 10 nucleotides. Several barcodes can correspond to a single sample, but each barcode is associated with only one sample. MGI Tech offers 96 variants of barcodes and a scheme for their optimal combination (Figure 1.).

**Figure 1.**
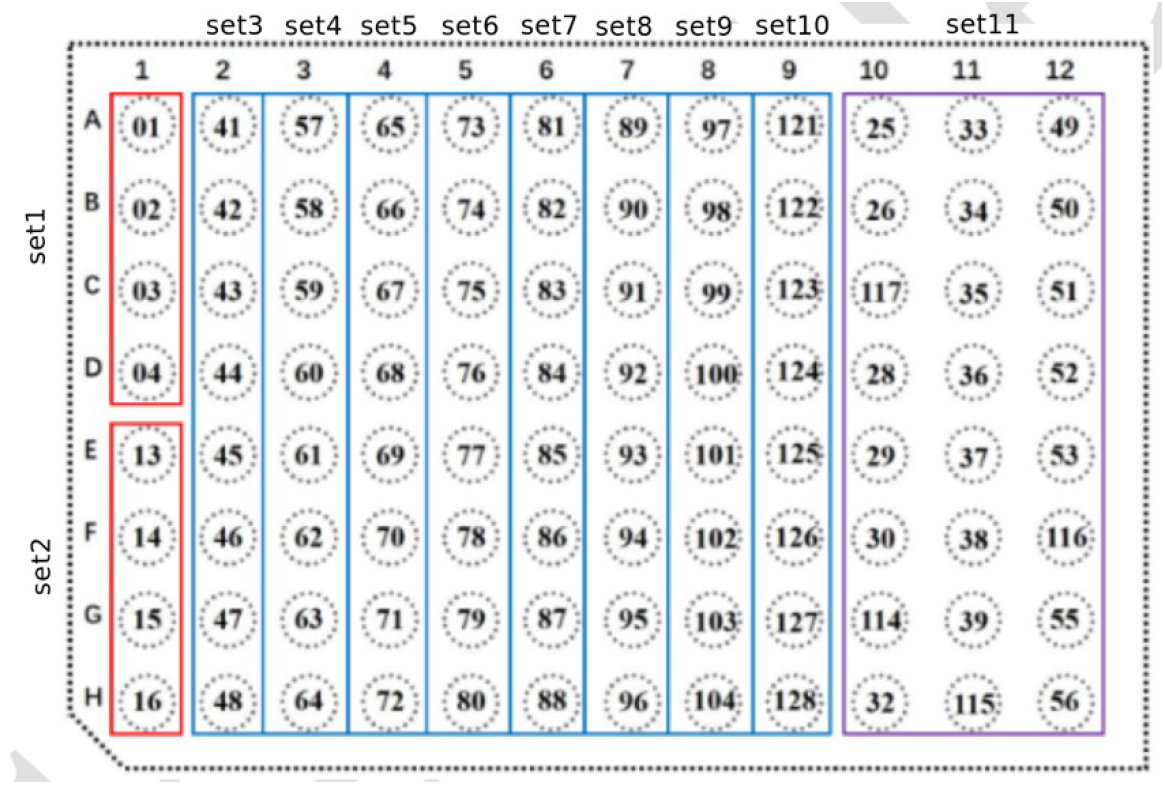
11 main compatible sets of barcodes according to the manual for MGISEQ-2000 [5]. Sets are highlighted with colored rectangles and signed around the perimeter.

This scheme of set combinations has a number of limitations: it is not possible to merge barcodes in different proportions; a certain number of samples are supposed to be used, which limits the opportunities of research. Thus, it can lead to uneven and inefficient use of barcodes. According to our subjective perception, MGISEQ-2000 is more sensitive to the correct balance of barcodes in a set than the Illumina HiSeq 2500 that we use as well.

To solve these problems and create alternative set variants we have designed the BC-store program. This tool combines the ability to analyze the alternative sets of barcodes with equal and custom ratios. The algorithm is based on analyzing the sensitivity of the MGI device to the concentration of nucleotides in the cycle phase at each of the positions in the barcode sequence. The method was used and tested in our previous MGISEQ-2000 runs. BC-store is an open-source software with source code freely available licensed under GPLv3. The BC-store is available on our lab’s website and GitHub [6, 7]. In this paper, we describe the development of the BC-store tool and its application in two real-world scenarios and in two user options from the command line and in the desktop version.

## Design and implementation

BC-store command-line version was developed in Python3 in order to be platform-independent; therefore, it can run under Linux, Mac, and Windows.

BC-store desktop version was developed in Python3 using QT-designer for desktop visualisation [https://doc.qt.io/qt-5/qtdesigner-manual.html] and works under Windows 10, which is installed by default on MGI Tech sequencers.

Users can collect sets of barcodes themselves, specify proportions and obtain results for matching barcodes, as well as add new barcodes in their own proportions. This format is particularly useful for workflows that require a lot of user interaction, such as selecting sets for sequencing.

## Formulation and verification of criteria

During sequencing in MGISEQ-2000, a complementary nucleotide is cyclically added to all nanoballs (DNB) on the flow cell, which means that each position on the barcode in all DNBs is read simultaneously. The nucleotide is determined by analyzing the measurements of the signal intensity from four fluorophores specific to each of the nucleotide types [8] in a specific sequencing cycle for a specific DNB. MGISEQ-2000 has limitations on the level of intensity perception (will be shown below on the example of launches). We defined two criteria – strong and lite; the first criterion (strong) was obtained based on the analysis of sets offered by MGI Tech. The second one (lite) was formulated by analyzing 20 runs of the MGISEQ-2000 sequencer in our laboratory. We found out that the MGI sensitivity is higher than the default set, so both the range by the number of barcodes and the intensity at each position can be increased.

To evaluate the success of sequencing each position, we used SNR values (signal- to-noise: shows the noise level for each of the four nucleotide types, calculated as the ratio of this nucleotide’s intensity to the intensities of the other three letters) (Figure 2) and FIT (essentially the same metric as SNR, but averaged over all 4 nucleotides), which are contained in html reports of the form “v300041900_run21_L02.summaryReport.html” for each lane and are generated by the device at the end of sequencing. Drops in FIT negatively affect the data quality, while a high level of FIT throughout the entire barcode reading indicates a better data quality (Figure 3). Next, it will be shown that drops in the FIT indicator occur at sites where the sets of barcodes are unbalanced.

**Figure 2.**
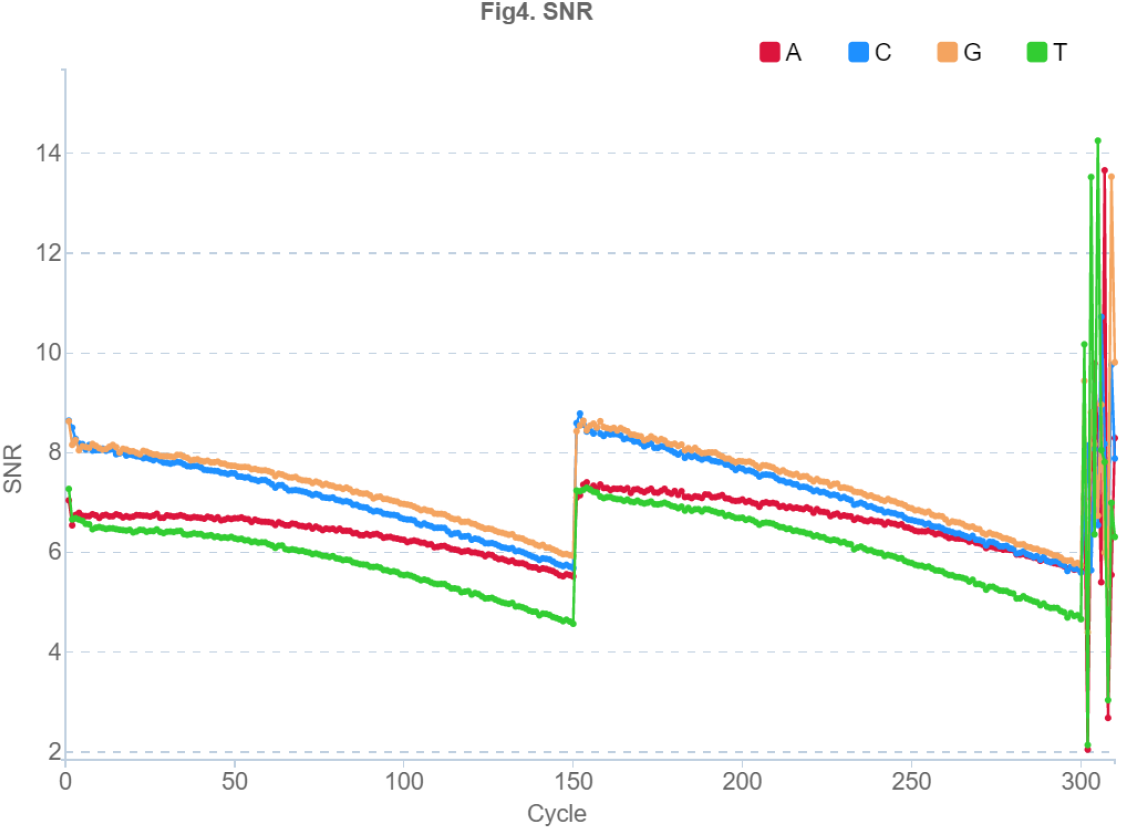
Example of a single lane SNR for a PE150 run on MGISEQ-2000. Sequencing cycles corresponding to the sequence number of the nucleotide in the reads are shown horizontally: 1:150 – forward read, 151:300 – reverse read, 301:310 – barcode sequence, SNR indicator is shown vertically. The indicator for each of the nucleotide types is marked by its own color (the colors are shown in the figure).

**Figure 3.**
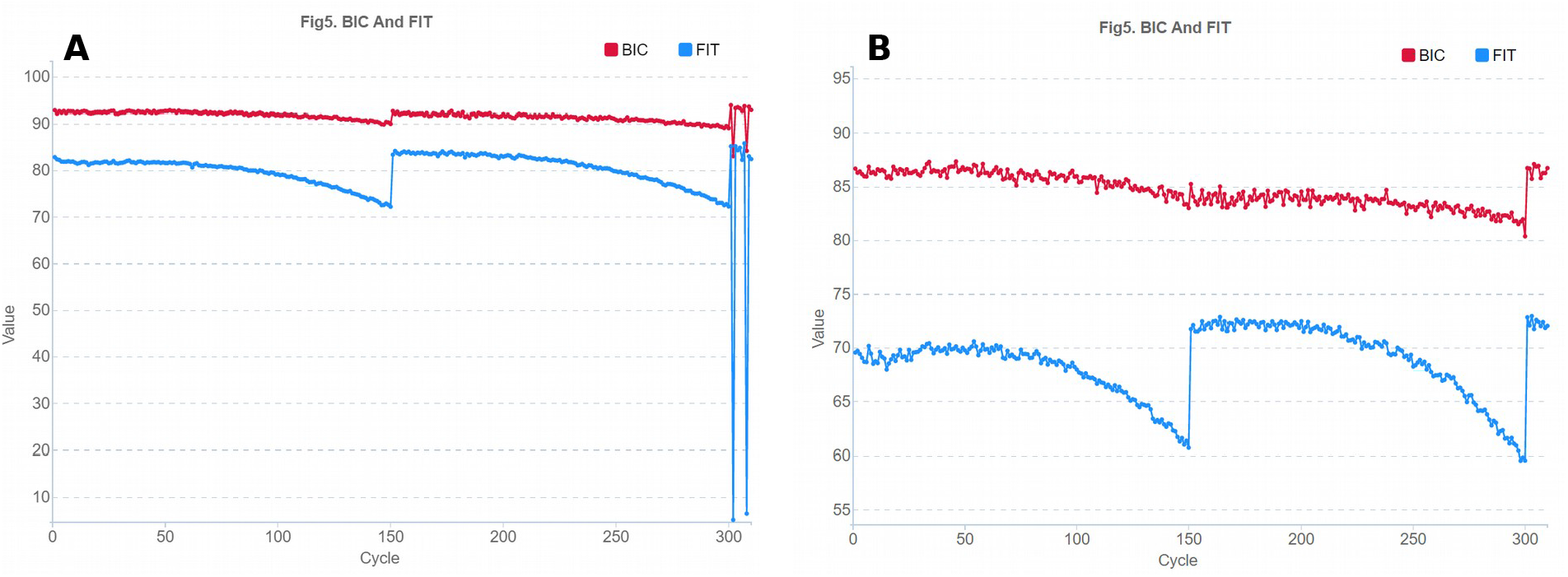
Example of a single lane FIT graph for a PE150 run on an MGISEQ-2000 with a non-optimal (A) and optimal (B) set of barcodes. Sequencing cycles corresponding to the sequence number of the nucleotide in the reads are shown horizontally: 1:150 – forward read, 151:300 – reverse read, 301:310 – bar code sequence, vertically – FIT (blue) and BIC (red) indicators which mean the probabilities or reliability of the base call results.

Notably, the cycle number increases with an increase in the error level (Figure 4). Since MGISEQ-2000 barcodes are always the last to be sequenced, it is important to minimize reading errors by balancing the set.

**Figure 4.**
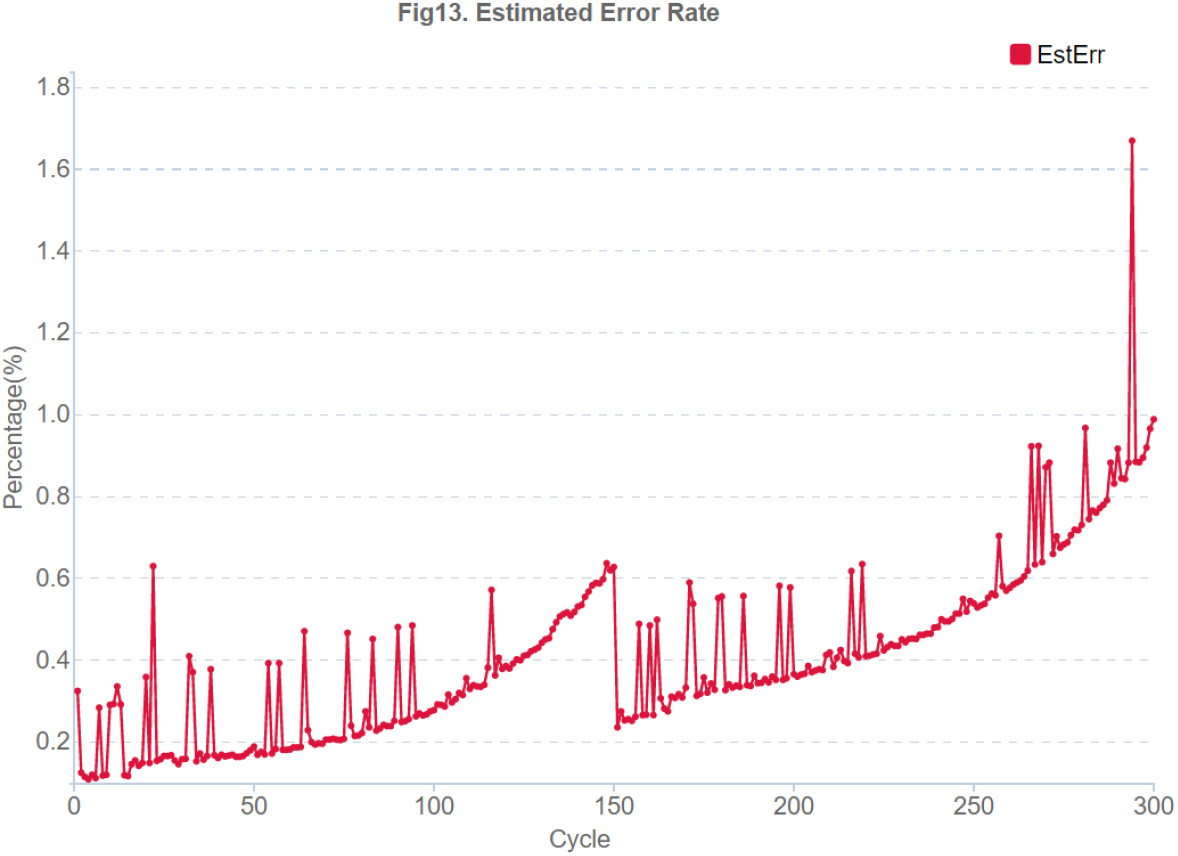
Error rate for a single-lane SNR for a PE150 run on MGISEQ-2000. The error rate increases with the increase in the cycle number.

We analyzed the output file SequenceStat.txt from the sequencer and concluded that MGISEQ-2000 can restore the original barcode sequence even if the individual letters are not read or read with an error (Figure 5 A).

**Figure 5.**
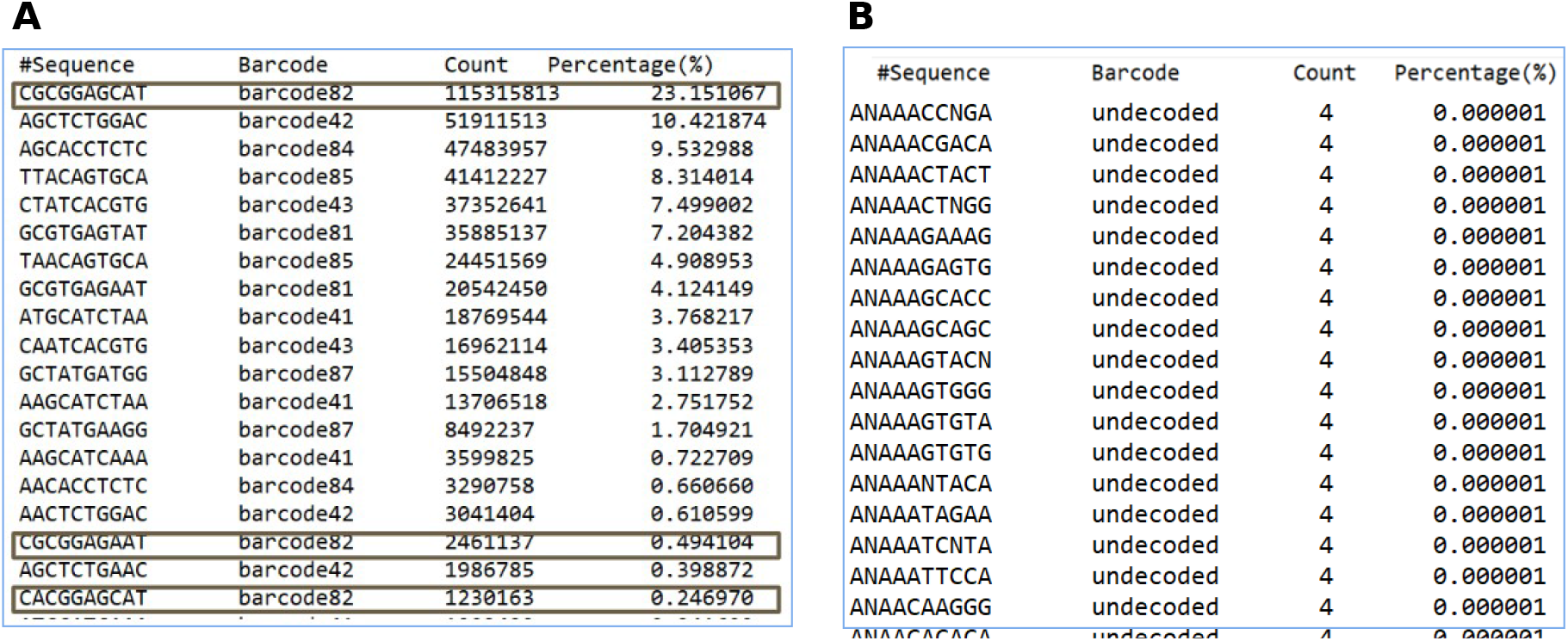
Example from the file SequenceStat.txt. A: barcode 82 was read in 23 percent of cases without error and in some cases with the recovery of the misread one nucleotide. B: read sequences of barcodes with more than two errors compared to the table of 96 sequences are assigned the status “undecoded”.

The algorithm uses the principle of comparing the found sequence of barcodes in a particular read with a table of 96 barcodes and searching for the closest one containing no more than 2 misread nucleotides. If the barcode is not found in the table, the read is assigned the “undecoded” class (Figure 5 B). FIT values below 10 indicate that the nucleotide was not recognized.

Based on the above, the cases with the presence of letters with a FIT below 10 can clearly lead to an increase in the share of undecoded nucleotides due to the inability to unambiguously restore the original sequence of the barcode.

FIT occurs when the sensitivity of the detector, which uses the signal intensity and the overlap of the glowing from neighboring DNBS to determine which nucleotide is at a given position, is limited. For example, if the diversity of each nucleotide type in neighboring DNBS is high, detection is successful (Figure 6 A). If one of the nucleotide types is overrepresented and the other one is underrepresented ( Figure 6 B), then the values of the SNR graph shift and, as a result, FIT drops in the corresponding positions of the barcode.

**Figure 6.**
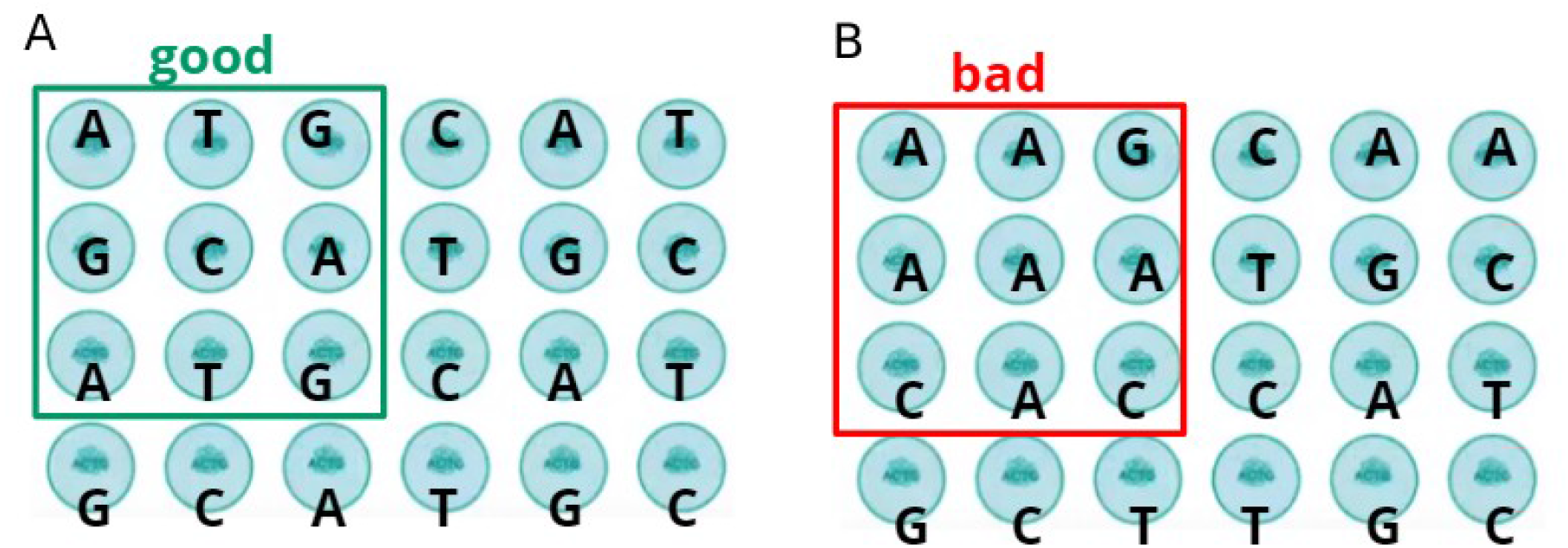
Schematic representation of the DNB on the flow cell. An ordered structure of DNB, each one is shown as a circle. The letter denotes the outer nucleotide on the nanoball to be identified in the current cycle. A 4– the level of radiation from each nucleotide type is approximately the same, the identification of nucleotides is successful, B – there is an overrepresentation of one of the nucleotide types and an underrepresentation of others, which leads to illumination and identification errors.

To test the hypothesis, we analyzed all sets offered by the manufacturer MGISEQ-2000 (Figure 1) for balance per each letter.

We wrote a script in Python3, that consists of two parts – computation and visualization. For sets 1-10, the fractions of each nucleotide at each bar code position were found to be equal to 0.25, and for set 11, they varied in a certain corridor (Figure 7). From the results obtained, conclusions were drawn about the limits of a strict criterion for selecting sets based on the NUCLEOTIDES FRACTION indicator.

**Figure 7.**
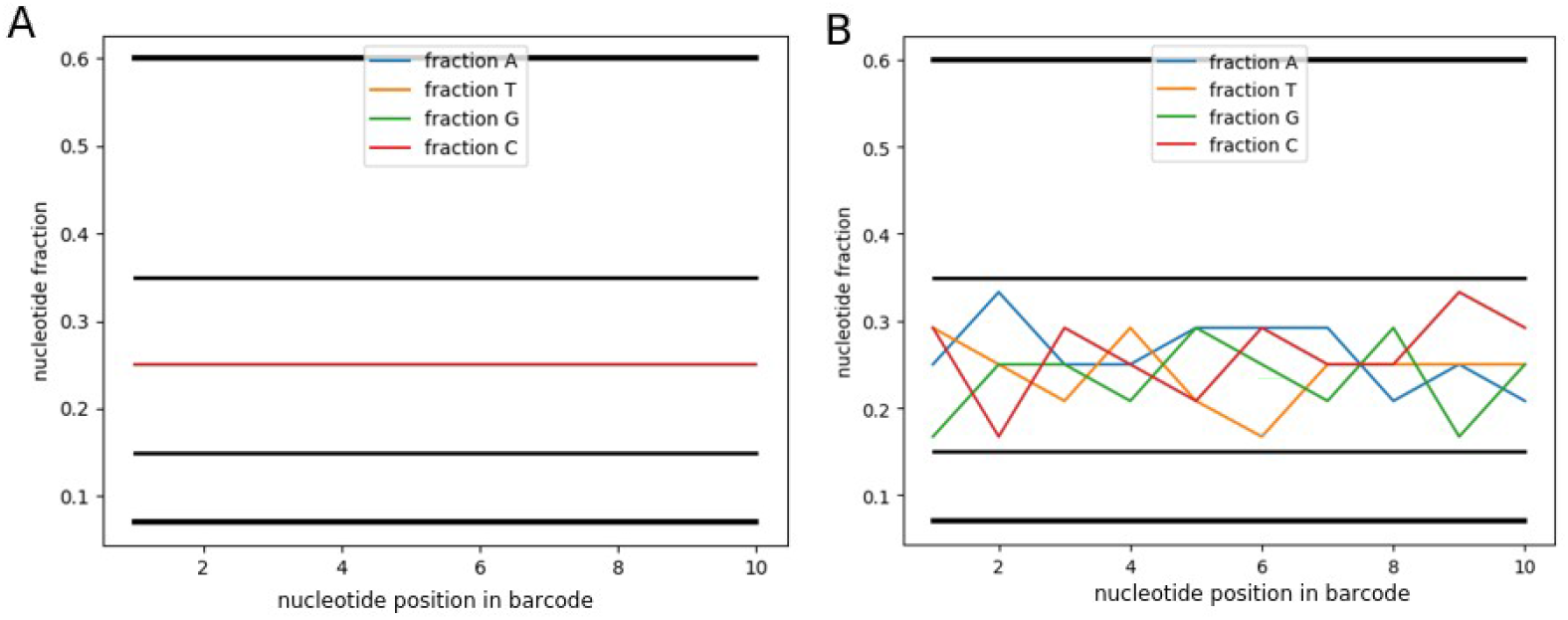
Example of the NUCLEOTIDES FRACTION script. A – for sets 1-10, the proportions of each nucleotide at each bar code position are equal to 0.25. B – for set 11, the proportions of each nucleotide vary within certain limits. Bold horizontal lines - lite criterion, thin lines - strong criterion.

An additional option was also introduced – mixing barcodes in different proportions. This option is relevant to the analysis of several samples from different sources, when it is difficult to observe equal concentrations of barcodes. For example, when one lane is loaded with samples for WGS x30 with significant differences in genome size. Problems with lowering FIT can be avoided if the concentration is taken into account when forming sets. The fact that the rate is worth considering can be clearly illustrated by changing the concentrations in set 1 from equal to unequal (Figure 8).

**Figure 8.**
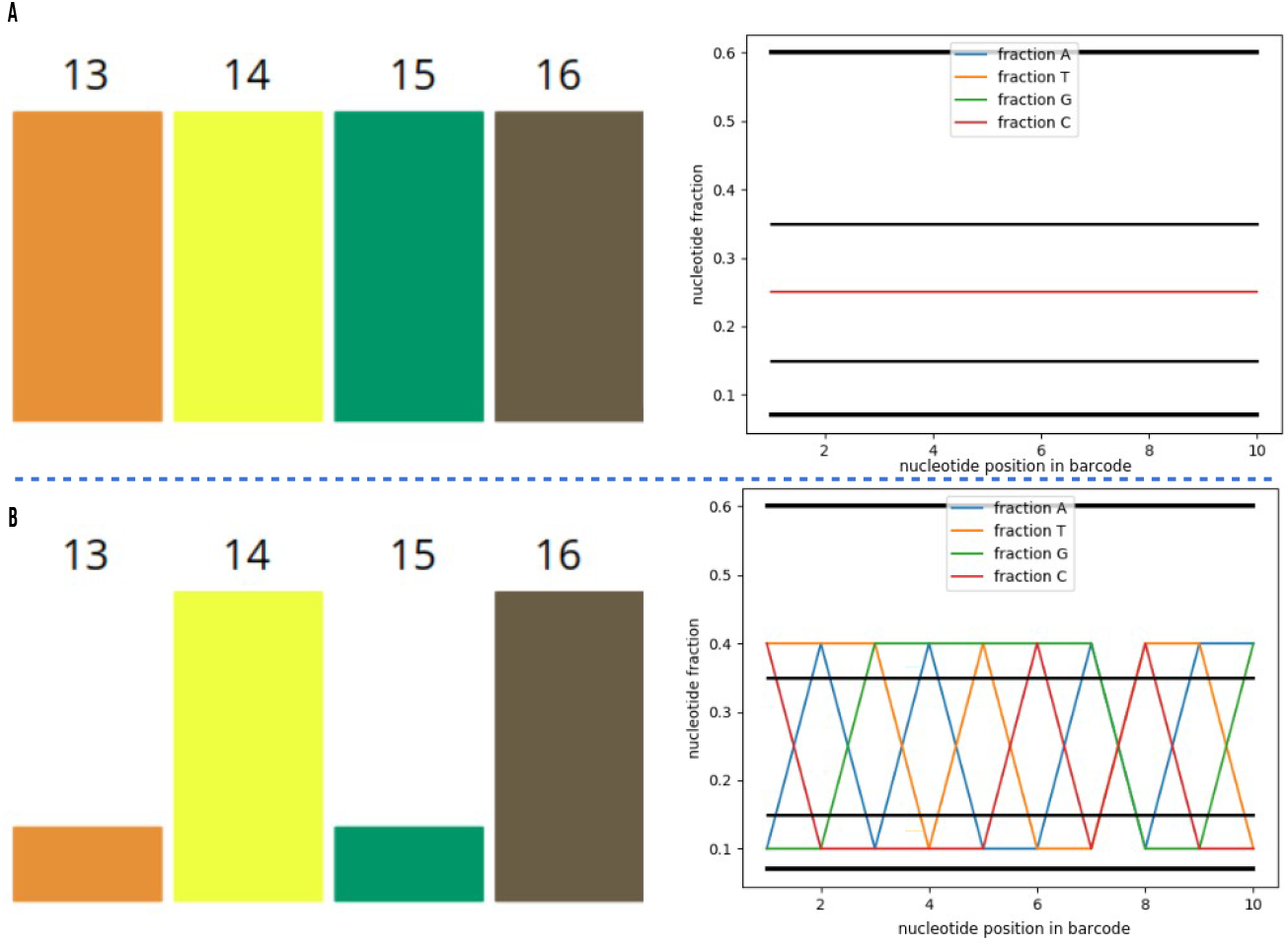
Changing the ratio in a set of barcodes. In the example of set 1, concentrations of 1:1:1:1 (A) changed to 1:4:1:4 (B) lead to an unbalance of the set from the initially perfectly balanced state.

Based on the analysis of 20 MGISEQ-2000 runs in our laboratory, the criterion was extended to the lite version, since in our runs the samples were often mixed in different proportions. For example, in the PE150 Z10 run, pooling samples with barcodes 87, 88, 89, 90, 91, 92, 93, 94, 95, 96 in proportion 9:9:9:9:5:5:5:5:5:5 did not lead to a drop in FIT (Figure 9 A). Only one position was within the strong criterion, 6 ones were on the border, and 3 ones were outside the criterion. In this case, the number of positions that fall outside the strong criterion (3) exceeded the algorithmically acceptable value (2). Therefore, the criterion can be extended to the following values ( Figure 9 B). An illustration that going beyond the lite limit leads to a drop in FIT is given for the Z2 and Z3 runs in Figure 10.

**Figure 9.**
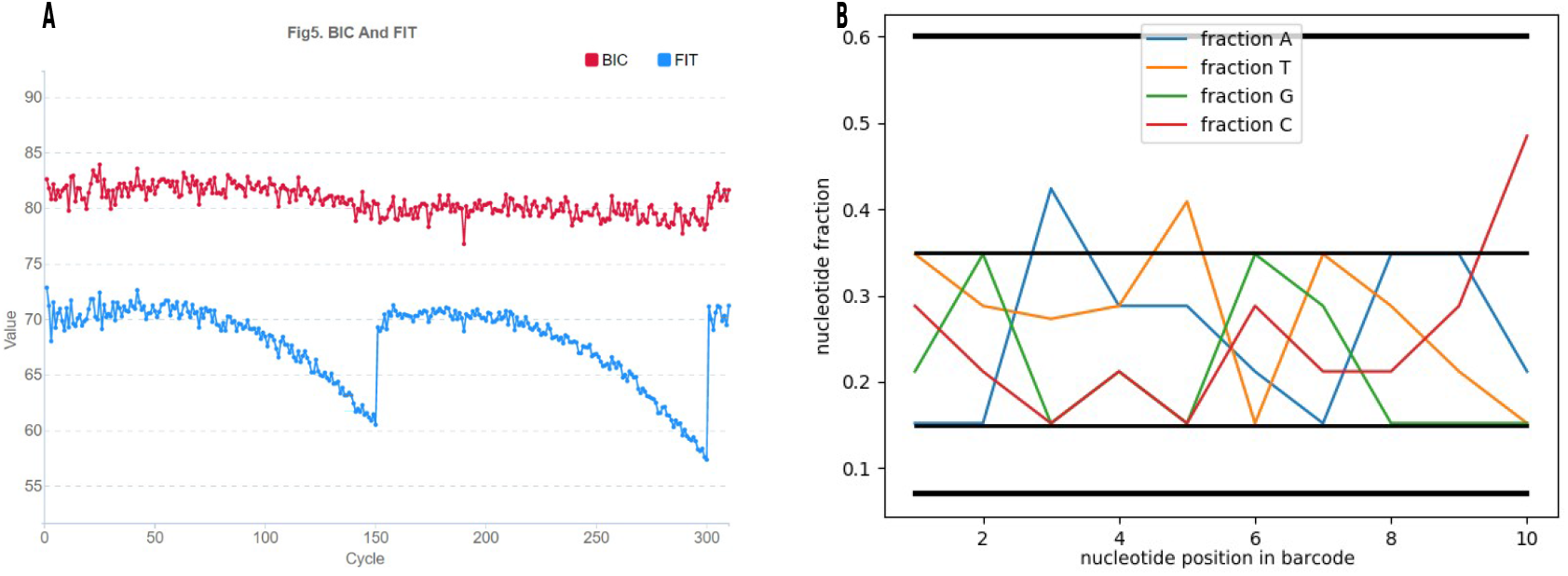
A – graph of the FIT and BIC distribution from the MGISEQ report for run Z10 and its NUCLEOTIDES FRACTION (B).

**Figure 1.**
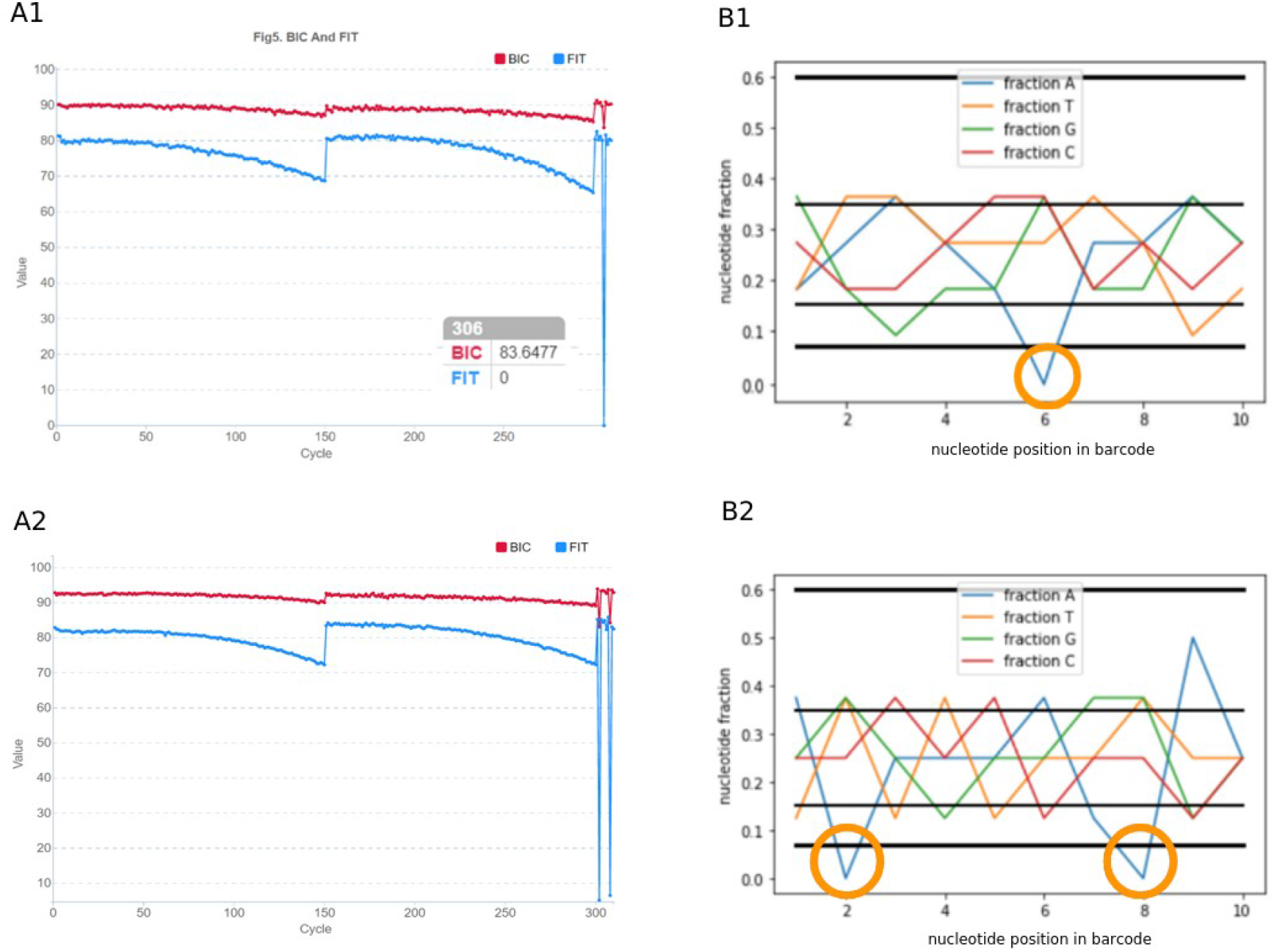
Examples of a direct relationship between FIT and lite criteria based on runs Z2 (A1, B1) Z3 (A2, B2): on the left, the graph from the MGI report, a drop in letters by FIT means that the nucleotide at this position was not recognized. On the right there is an example of how the script works, the drop outside the criteria is highlighted with circles. The FIT drop and the criterion drop are in the same positions in the bar code.

We also analyzed the number of nucleotides making the barcodes different. A script was written to compare the sequences of barcodes in pairs (Supplementary). The minimum difference is 4 nucleotides, while the largest one is 10. Obviously, in order to avoid “merging” of barcodes in a set, the number of mistakenly read nucleotides during sequencing should not be equal to or exceed the smallest number of differences between barcodes from the set among all those obtained during pairwise comparison. In general, the number of errors during their sequencing should not be equal to or exceed 4.

To check the decrease in the share of undecoded data generated by ZebraCall in the case of more than 2 unrecognized nucleotides in the barcode reading, we used the data recovery script (https://github.com/gateswell/SplitBarcode). In one of our runs in PE150 mode on a single lane, we received 2.4 GB and 2.5 GB for forward and reverse reads of undecoded files, respectively, comprising no more than 2% of the total data from this lane – this is the usual proportion for a well-passed run on MGISEQ-2000. We varied the parameters of the number of mismatches (2, 3, and 4) from the list of barcodes with which the sequenced barcodes are compared (17 pieces used in the launch or all 128 barcodes), and determined how much the undecoded percentage decreases after applying the script (Table 1). The size of the undecoded file for forward and reverse reads was found to be equal to those obtained through the algorithm if we took the mismatch value equal to 2. In the case of a mismatch equal to 3 and 4, the undecoded number was significantly reduced. Reducing the list of barcodes that the sequenced barcodes were compared did not significantly reduce the size of the undecoded fraction.

**Table 1.** The impact of the number of allowed mismatches in the bar code on the proportion of saved data from undecoded

| Barcodes in list | mismatches | undecoded R1, byte | undecoded R2, byte | % rescued data |
| --- | --- | --- | --- | --- |
| 17 | 2 | 2502659710 | 2652072230 | -0,47 % |
| 17 | 3 | 2374062925 | 2515798025 | -5,59 % |
| 17 | 4 | 1746296040 | 1850552520 | -30,55 % |
| All 128 | 2 | 2514592410 | 2664717330 | 0 % |

We also analyzed the Zebracall application installed on MGISEQ-2000. It converts .cal files to fastq ones. There are several launch options in the app folder: the first one starts automatically when sequencing is complete. The second one (C\:ZebraCallV2\ client.exe) allows the user to run Zebracall independently with the required parameters. By default, the number of mismatches is also set to 2. Increasing the possible number of mismatches significantly reduces the value of undecoded nucleotides. However, we do not recommend using a value exceeding 2, since in addition to the unbalanced set, there are other reasons that cause non-reading of letters during sequencing. The latter can lead to “connecting” samples with similar barcodes, for example, MGI Tech has barcodes that differ only by 4 letters out of 10.

Thus, we have demonstrated that the most important criterion determining the quality of sequencing data is the balance of barcodes in the set. It is also possible to have an unbalance for no more than two positions. We will further describe how BC-store works and how results are interpreted. Since we cannot guarantee that in addition to several mismatches, there will be no additional reasons for non-reading of nucleotides at launch, we do not allow exceeding the criteria for any bar code position by default. This approach yields good results in practical work, yet users can act at their own discretion. In any case, BC-store will be a useful application for selecting barcodes.

## Results

### Architecture

BC-store is available in two versions: desktop for Windows and command line for operating systems with pre-installed Python3, available for download at [6, 7].

### How it works from the command line

The program is launched by running it from the command line:

~~~
python3 [path to bc-store_script.py] [command] [options by
whitespace]
~~~

<u>There are 4 main commands:</u>

`check_set` – checks the current set based on the proportions according to the selected criterion

`add_to_set` – selects the desired number of barcodes and adds to the existing set, taking into account the proportions according to the selected criterion from the selected set of barcodes

`help` – help with the examples of commands and explanation of options and format for entering parameters

`MGI_sets` – displays successful sets of barcodes according to the MGI manual

After running the `check_set` commands and if the set selection is successful, the user will be provided with a graph by the `add_to_set` command. To run a new command, you need to close the drawing and save it if necessary. To use the script, some standard packages specified at the beginning of the python3 file are required. Examples of how BC-store works from the command line in the case of equal and unequal proportions of barcodes mixing in a set are shown in Figure 11. The first one is activated by the command:

~~~
python3 bc-store_script.py check_set set=13,14,15,16
rate=1,1,1,1 criteria=strong
~~~

The second one is activated by the command:

~~~
python3 bc-store_script.py check_set set=13,14,15,16
rate=1,4,1,4 criteria=strong
~~~

**Figure 11.**
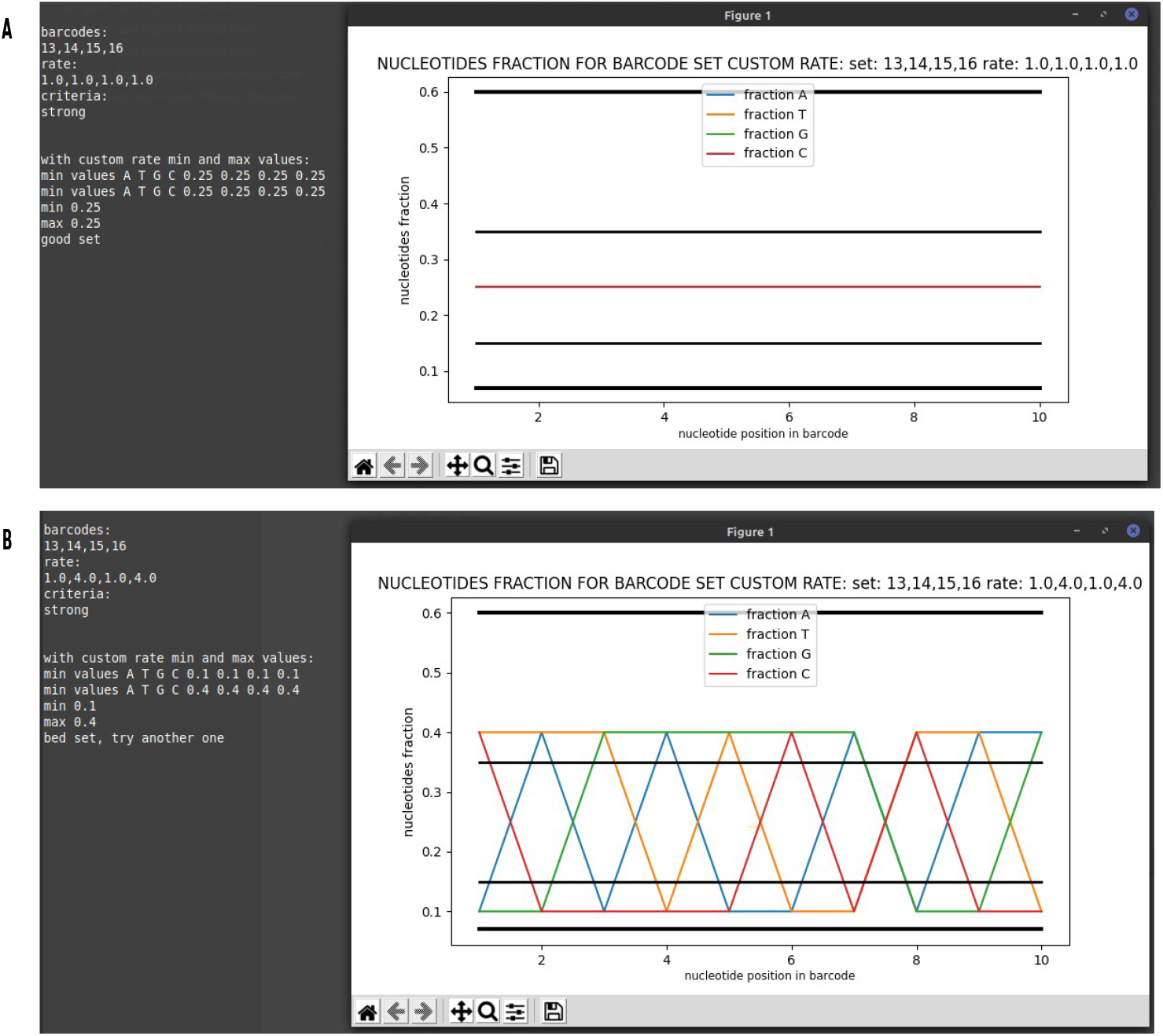
Example of BC-store operation from the command line in case of equal (A) and unequal (B) proportions of barcodes mixed in a set.

### How it works on the desktop version

The desktop version with the numbering of input fields and buttons is illustrated in Figure 12. When you start the desktop version, an additional window opens where errors are displayed so that you do not need to close it for the program to work correctly. The main areas of work are set analysis (sections on Figure 12 : 1, 2, 3), set selection (sections on Figure 12: 1, 5, 6, 7, 9, 10) and output of basic sets (section on Figure 12: 8). After clicking the “Click to analyse set” button and, if there are additional set barcodes, when clicking the “Click to analyze adding” button, the user will be provided with a graph. Do not press buttons 3 and 10 until you have filled in all the required fields. The program presents the output information on user-defined barcodes, mixing proportions, values of nucleotide representation, and the main information on whether this set meets the strong and lite criteria (Figure 13).

**Figure 12.**
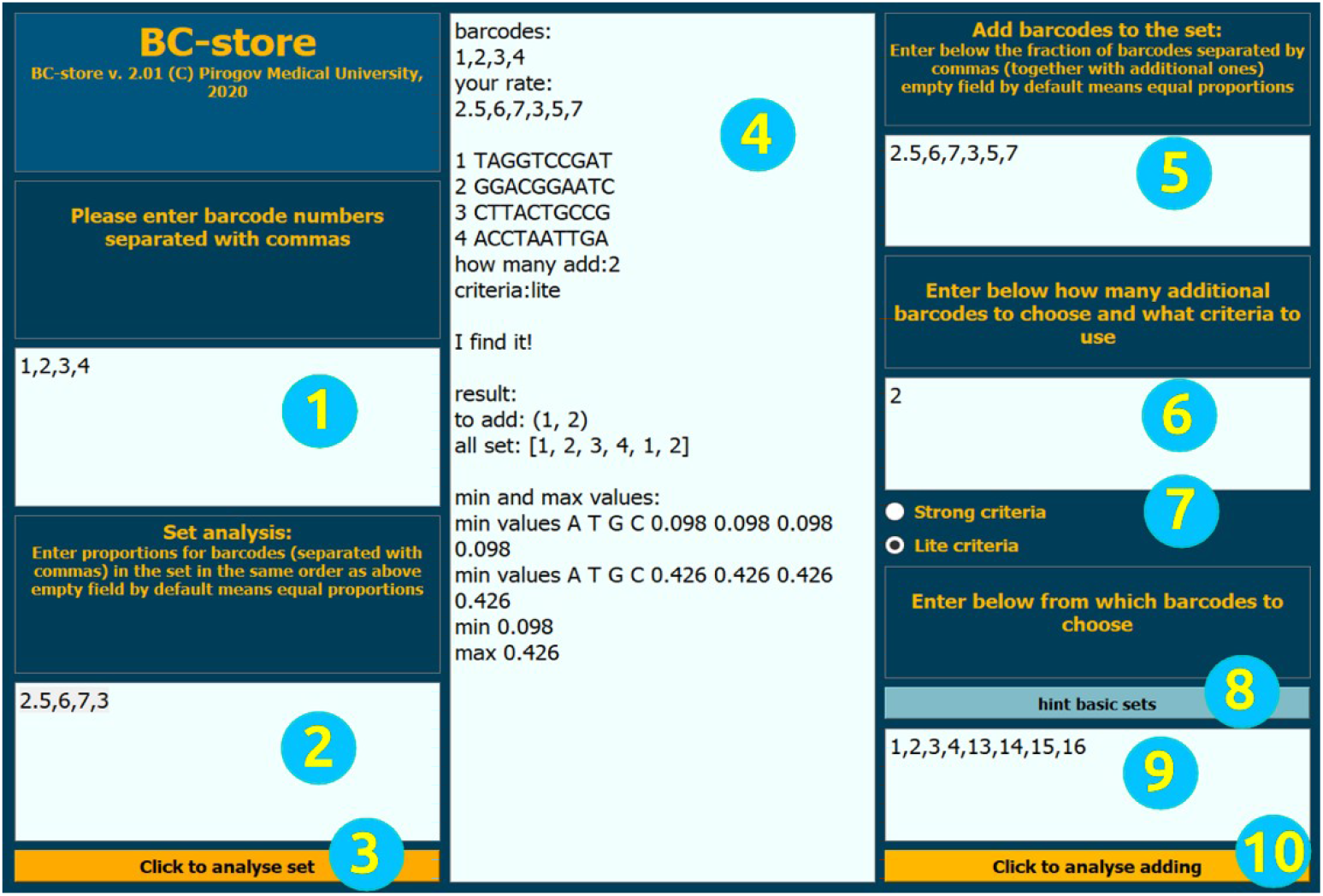
Layout of the desktop version with numbering of input fields and buttons. 1 – enter the current set, 2 – enter set proportions, 3 – set analysis button, 4 – output field, 5 – input proportions of the current set with the new barcodes, 6 – enter the number of new barcodes, 7 – select criteria, 8 – tip for base sets, 9 – input the barcode to choose from, 10 – start analysis of set selection.

**Figure 13.**
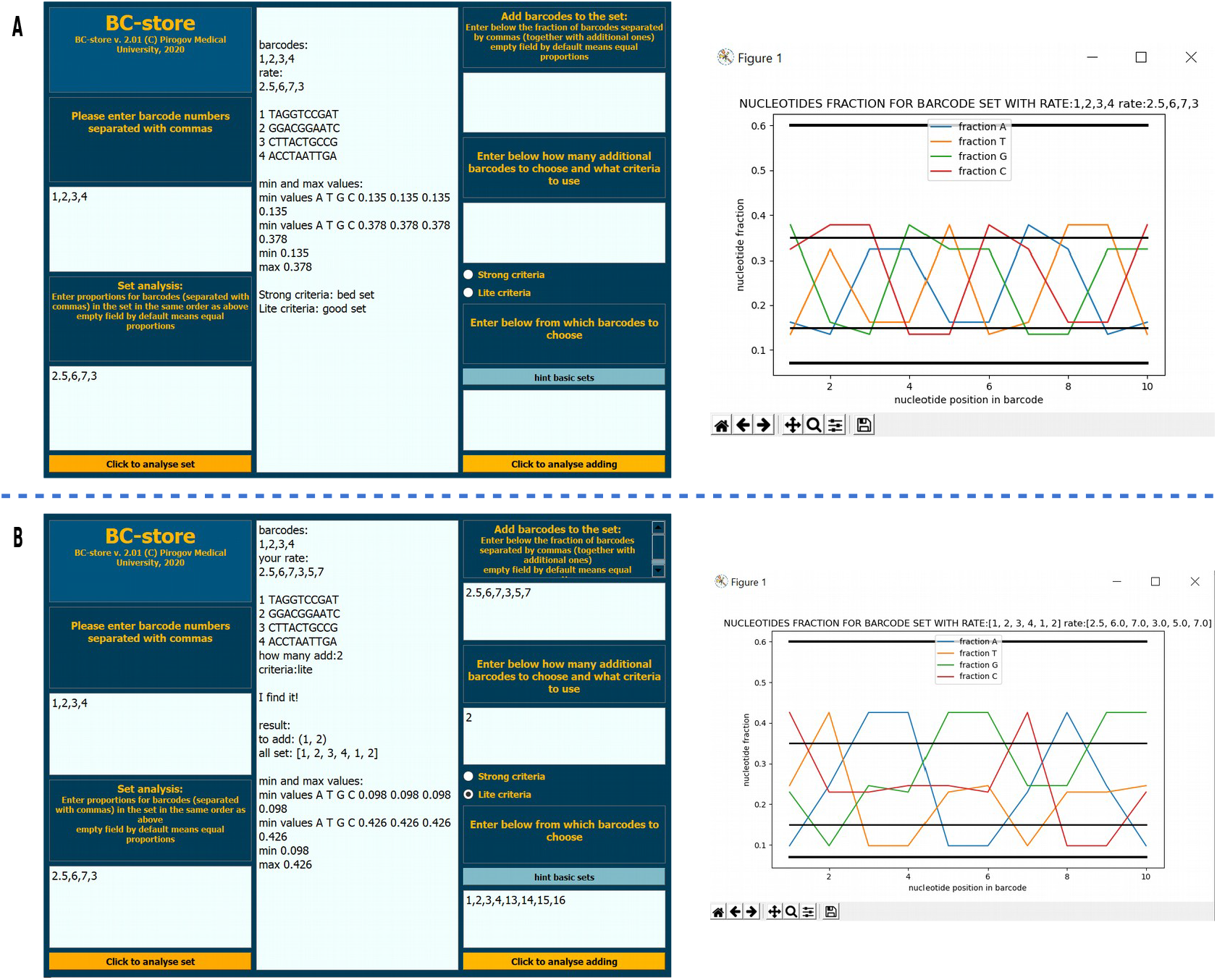
A – an example of how BC-store works in case of equal proportions of barcodes in a set. In the case of equal proportions, the field for entering shares should be left free. B – an example of how BC-store works if the other barcodes are added to the set.

### Recommendations for MGI base sets (Figure 2)

Each of sets 1-10 is perfectly balanced when used in equal proportions, so it can be added to any ready-made set, equal proportions and a full set of barcodes should be observed. Furthermore, these sets can be combined with each other at any ratio, equal proportions within each set should be maintained.

## Discussion

To demonstrate the effective operation of the BC-store on MGISEQ-2000 here we discuss the results for several launches. In particular, we used the non-standard combinations of barcodes that can be considered successful and unsuccessful, as well as the values of the undecoded data proportion, the number of dropped nucleotides in the FIT barcodes. Characteristics of runs and BC-store results for some of our PE150 runs on MGISEQ-2000 are shown in Table 2 and Figure 14. Thus, BC-store is firmly established in the routine practice of our laboratory and allows us to successfully combine samples in a variety of situations.

**Table 2.** Characteristics of PE150 mode runs on MGISEQ-2000 with good and bad quality.

| Run | Set of barcodes | Rate | Number of drops in FIT | % undecoded | Graphs in figure 14 |
| --- | --- | --- | --- | --- | --- |
| **Z5** | 35, 52, 73, 87, 88 | equal | 9 | **22** | A |
| **Z6** | 44, 81, 100 | **90, 12, 12** | 10 | **63** | B |
| **Z7** | 44, 45, 59, 60, 61, 62, 65, 66<br>67, 68, 69, 70, 71, 72, 73, 74, 83 | equal | 0 | **0** | C |
| **Z8** | 41, 42, 43, 44, 46, 47, 48, 61, 62, 63,<br>64, 65, 66, 67, 68, 69, 70, 72, 73, 74,<br>75, 76, 77, 78, 79, 80, 81, 82, 85, 86,<br>87, 88, 89, 90, 91, 99 | equal | 0 | **1.7** | D |
| **Z9** | 28, 29, 30, 32, 34, 37, 55, 57, 58, 59,<br>60, 61, 62, 63, 64, 65, 66, 67, 68, 69,<br>70, 71, 72, 73, 74, 75, 76, 77, 78, 79,<br>80, 81, 82, 83, 84, 85, 86, 87, 88, 117 | equal | 0 | **2** | E |
| **Z10** | 87, 88, 89, 90, 91, 92, 93, 94, 95, 96 | **90, 90, 90, 90,**<br>**50, 50, 50, 50,**<br>**50, 50** | 0 | **1.75** | F |

**Figure 14.**
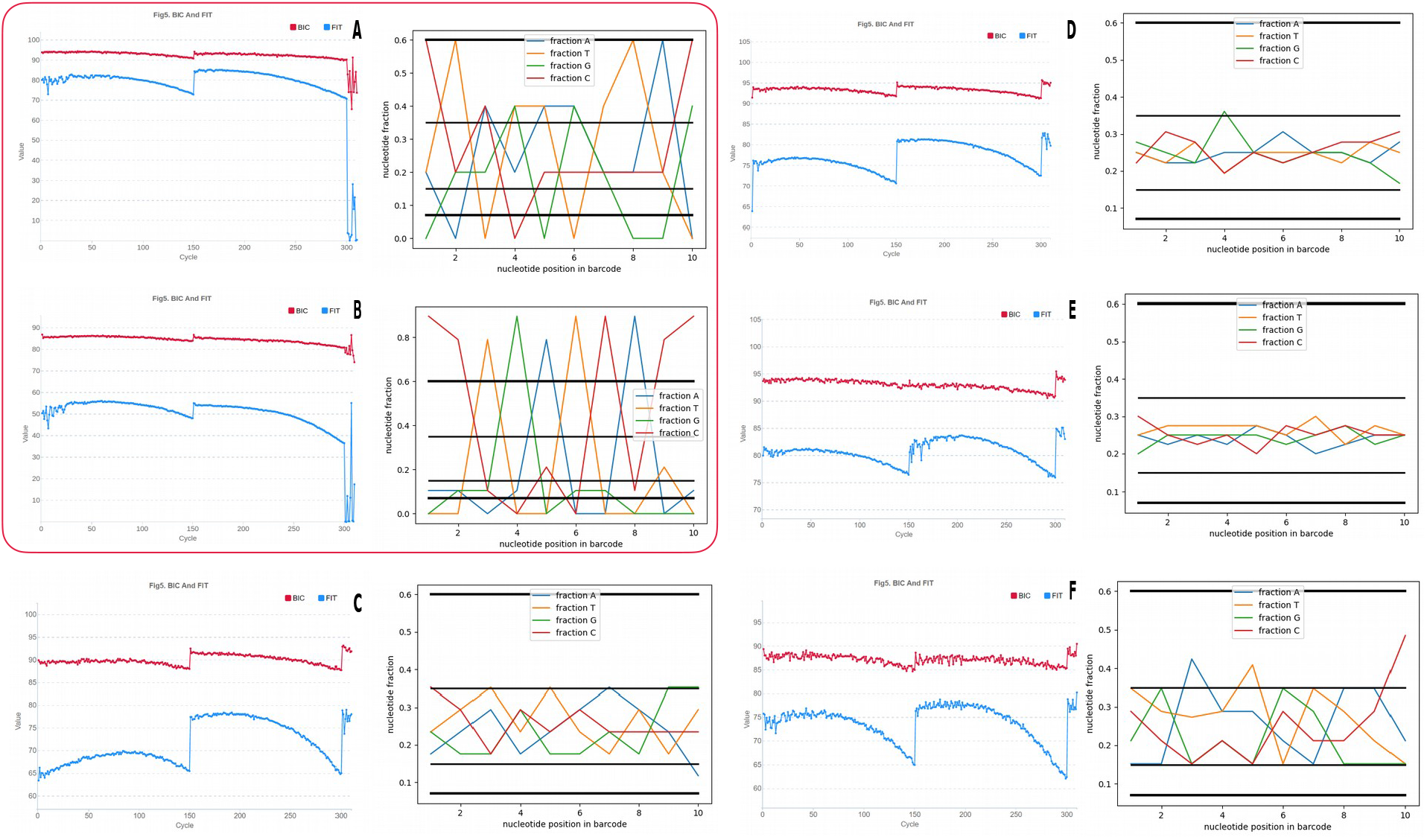
FIT graphs and BC-store results for analyzing barcode sets for unsuccessful (A, B) and successful (C-F) launches in our laboratory in PE150 mode on MGISEQ-2000.

## Availability and future directions

The main website for BC-store is https://store.genomecenter.ru/bc-store where the program itself, links to Github command line version, tutorials and documentation can be found.

## Funding

This research was funded by grant №075-15-2019-1789 from the Ministry of Science and Higher Education of the Russian Federation allocated to the Center for Precision Genome Editing and Genetic Technologies for Biomedicine

## Conflicts of Interest

The authors declare no conflict of interest.

## Notes

### Competing Interest Statement

The authors have declared no competing interest.

