## Supplementary figures and images for "BC-store: a program for mgiseq barcode sets analysis"

### sup_A

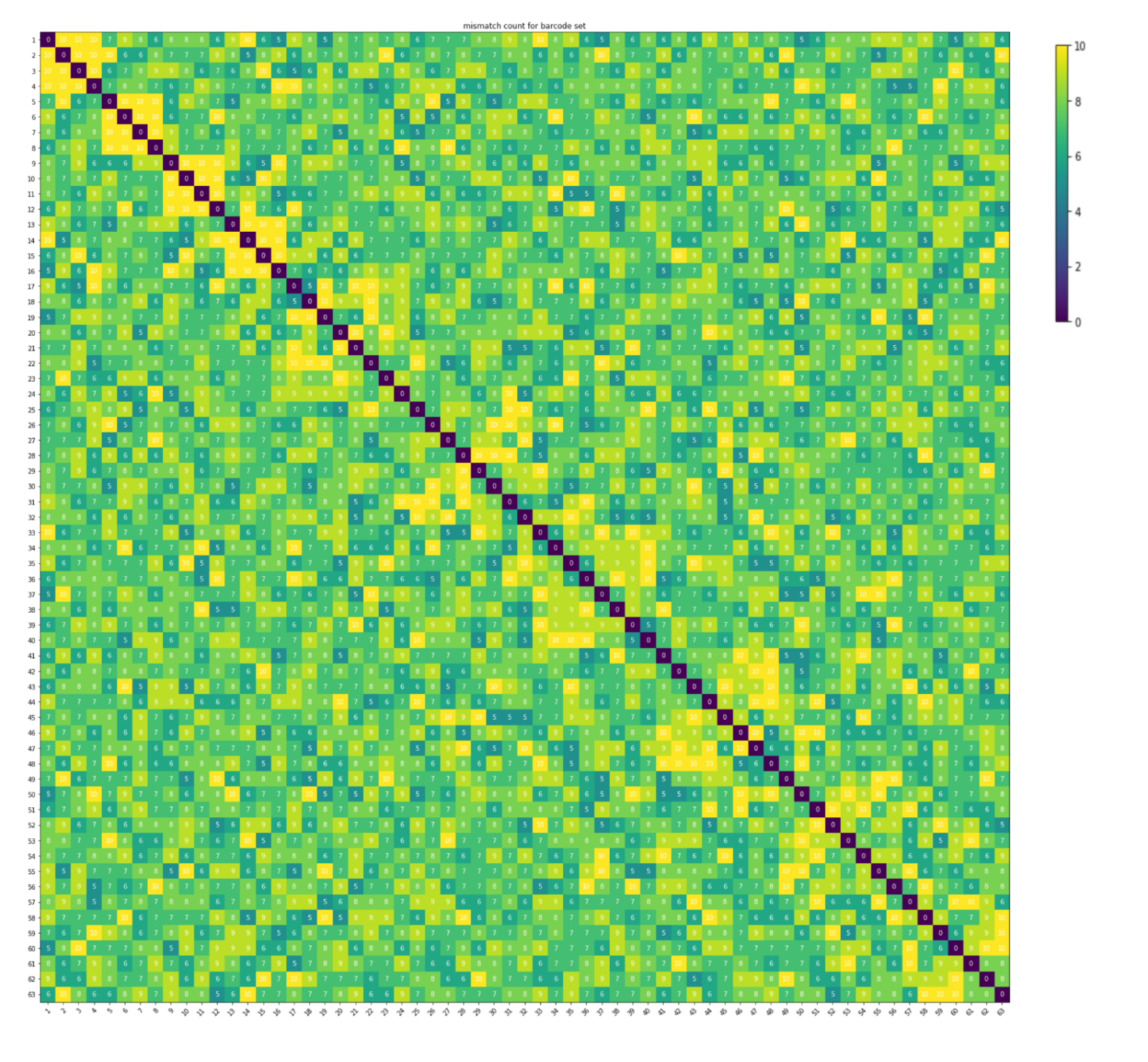

### sup_B

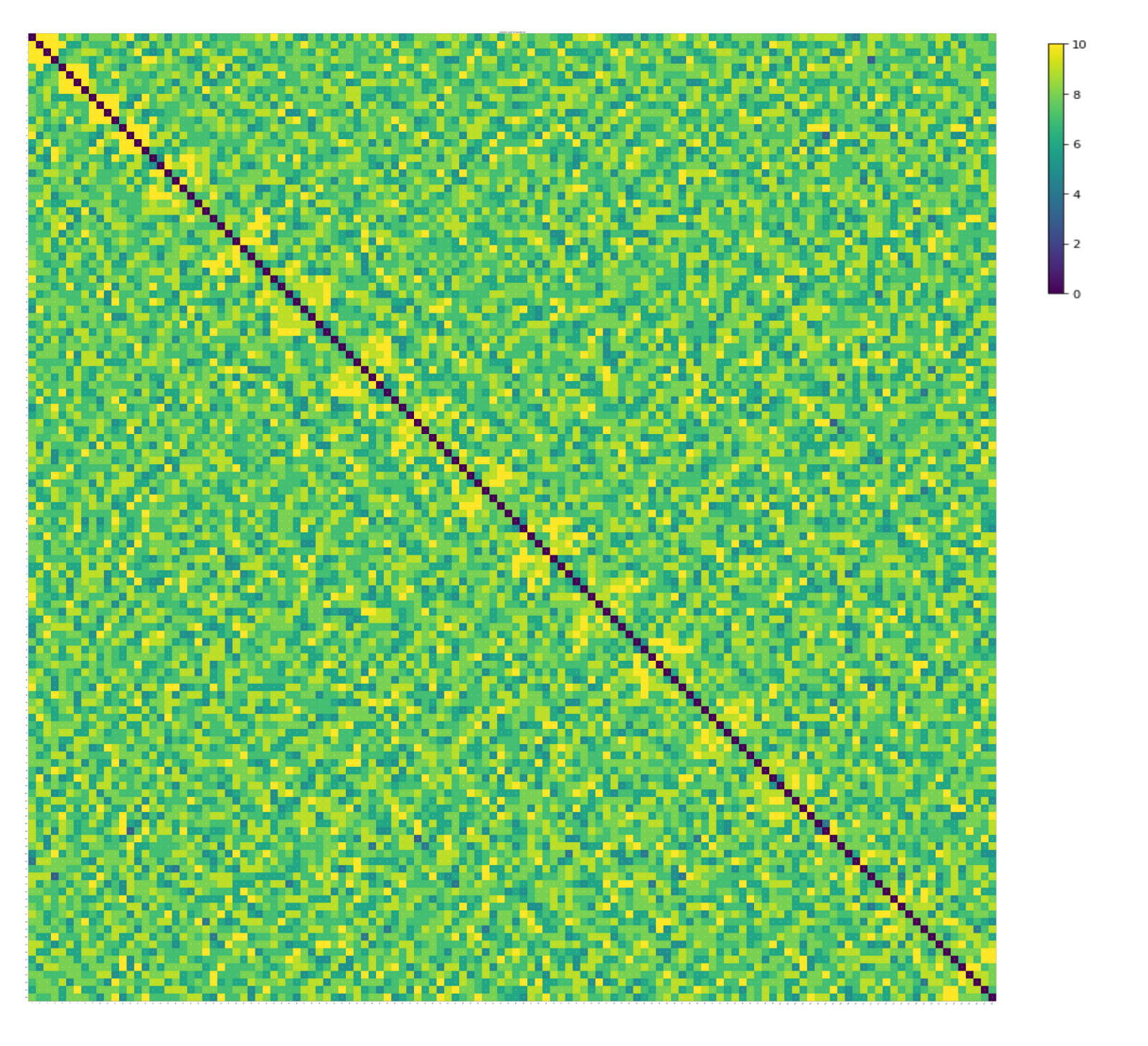
